# The Cell Subtypes Selection by Genes (CSSG) algorithm for discovering cell populations in high resolution

**DOI:** 10.1101/2023.04.20.537649

**Authors:** Jakub Kubiś, Maciej Figiel

## Abstract

The recent massive improvements in transcriptomics and single-cell technologies have led to a rising volume of data and demand for advances in bioinformatics processing. Existing methods are not fully capable of discovering genetic markers responsible for high-resolution cellular tissue heterogeneity, cell lineages during organism development, and cell differentiation with rare intermediate populations. In response to demand, we have generated a new Cell Subtypes Selection by Genes (CSSG) algorithm which is supported by a dedicated and fully automatic JSEQ^®^ pipeline. The new CSSG algorithm is iterative, parallel, and able to make decisions for discovering cell populations in tissues based on transcript occurrence in cells. The CSSG/JSEQ is complemented by a new strategy and specialized algorithm for the naming of cell populations. Our approach allows for high-resolution tracing of cell populations, finding relations and hierarchy between them, particularly important for complex tissues such as the brain. The pipeline allows the establishment of developmental, differentiation, and pathogenic trajectory and takes a “snapshot” of a current physiological or pathological cellular stage of the investigated organ at the transcriptional level.

## Introduction

Single-cell RNA sequencing (scRNAseq) generates high-volume of transcriptomic data sets containing information about each cell’s gene expression separately, demanding complicated multi-step bioinformatics analysis. The required approach is a reduction of dimensionality (expression of genes and their relations) by specialized methods (PCA, UMAP, or tSNE) to be able to percept and interpret the data (1). Reduced dimensions permit further clustering analysis, which allows the exploration of the primary sources of heterogeneity of cell composition in the tissue. Such an approach is prone to the loss of biologically significant information. In the case of scRNAseq, it can cause a loss of information about rare cell populations that are normally neglected but exhibit important biological functions, such as in the brain. In addition, disease development may start from a small change in a small population of cells, of which recognition is crucial for developing therapy. Therefore, we decided to create the single-cell algorithm called Cell Subtypes Selection by Gene (CSSG^®^) methods supported by a fully algorithmic-based and automatic JSEQ^®^ pipeline with a custom-developed cell population naming algorithm. The CSSG algorithm conducts the iterative summing of matrices containing information on gene occurrence in cells belonging to one cluster, which is subsetted into detailed cell subtypes. Based on algorithmic decision-making, the CSSG algorithm delivers information about cell intermediate stages, maturation status, rare population functions, trajectories, and interactions between them.

The brain is a very complex structure where a variety of cell types express a unique set of genes at different steps of development and maturation (2). These cellular populations and universal gene sets are essential for maintaining brain regions’ general anatomical positioning and connectivity between regions to preserve the essential state of brain functionality required. Therefore, a slight difference in cellular populations may greatly impact brain structure, functionality, and neurodegenerative disease development, e.g., Alzheimer’s or Parkinson’s.

Therefore the cell heterogeneity of the brain is an excellent model to test our CSSG algorithm and to unravel the high-resolution cell complexity of the organ. Using our CSSG/JSEQ, we performed deep bioinformatics analysis of the brain single-cell / single-nuclei data, including 812945 cells. We revealed 620 subtypes of various brain cell populations from different brain regions.

Three-step pipeline validation was conducted by comparing the cell subtype composition of clusters with and without the CSSG, determining the trajectories of oligodendrocyte development, and finally by tracking the relation of the cell populations possibly related to Alzheimer’s disease with specific expression of AD genes such as Apoe, App, Mapt, Psen1, Psen2, and Trem2. We concluded that the CSSG/JSEQ approach is suitable for complex analysis of scRNAseq data to reveal the very high resolution of rare cell populations, which are highly significant in brain development and disease.

## Materials and methods

### Data

The single-cell raw count sequencing data were acquired from public repositories. The data were used for pipeline functional tests, high-resolution cell type discovery, and analysis of cell types causative to Alzheimer’s disease. The data for the mouse hypothalamus, mouse thalamus, and human organoids were acquired from NCBI GEO. Mouse hypothalamus [HYP] (7 329 cells) data contain lateral hypothalamic areas from male and female mice, GSM3562050 and GSM3562051, respectively (3). Mouse thalamus [TH] (7 365 cells) from the embryo (E12.5): GSE122012 (4). Three and six months, human brain organoids [HBO] (38 987 cells) were acquired from GSE129519 (5). Data from the adult mouse for motor cortex [MCTX] (13 539 cells) (MD704, MD705), auditory cortex [ACTX] (35 594 cells) (MD717, MD720, MD21), orbitofrontal cortex [OCTX] (19 531 cells) (MD716, MD717), entorhinal cortex [ECTX] (80 653 cells) (MD720, MD721, MD698699), hippocampus [HIP] (6 555 cells) (MD718, MD719), and cerebellum [CER] (611 034 cells) (6) were acquired from Broad Institute data portal https://singlecell.broadinstitute.org/. The data used for the analysis were derived from different sequencing methods, including single-cell and single-nucleus. Therefore for unification purposes, all mitochondrial genes and metabolism-related genes were removed based on genes from MitoCarta2.0 (7). The data processed by the pipeline are available on NCBI GEO.

All computation stages were performed on a workstation configured with AMD Ryzen Threadripper 3960X 24-Core 3.80 GHz processor, 256GB of RAM, Windows 10 Pro running Ubuntu 20.04 LTS virtual machine. Automatical analysis: JSEQ_scRNAseq (included software and their versions on GitHub in documentation). Manual analysis RStudio Version 1.3.1093 with R version 4.0.3.

## JSEQ^®^ single-cell analysis pipeline

All analyses of raw single-cell data were performed using our JSEQ^®^ single-cell analysis pipeline (Fig. 1A). The pipeline scripts and the source code is available on GitHub [https://github.com/jkubis96/JSEQ_scRNAseq]. The pipeline and its source code are available for academic use. The JSEQ^®^ pipeline contains modules for all applications arranged and controlled by the queue and control points that prove that all conditions required for the next step are met. This way, interoperability between consecutive steps/points is established based on storing temporary data in the project working directory. Moreover, the data for the pipeline can be delivered in multiple formats. The pipeline allows for analyzing both single-cell or single-nuclei data in different input formats.

**Figure 1.**
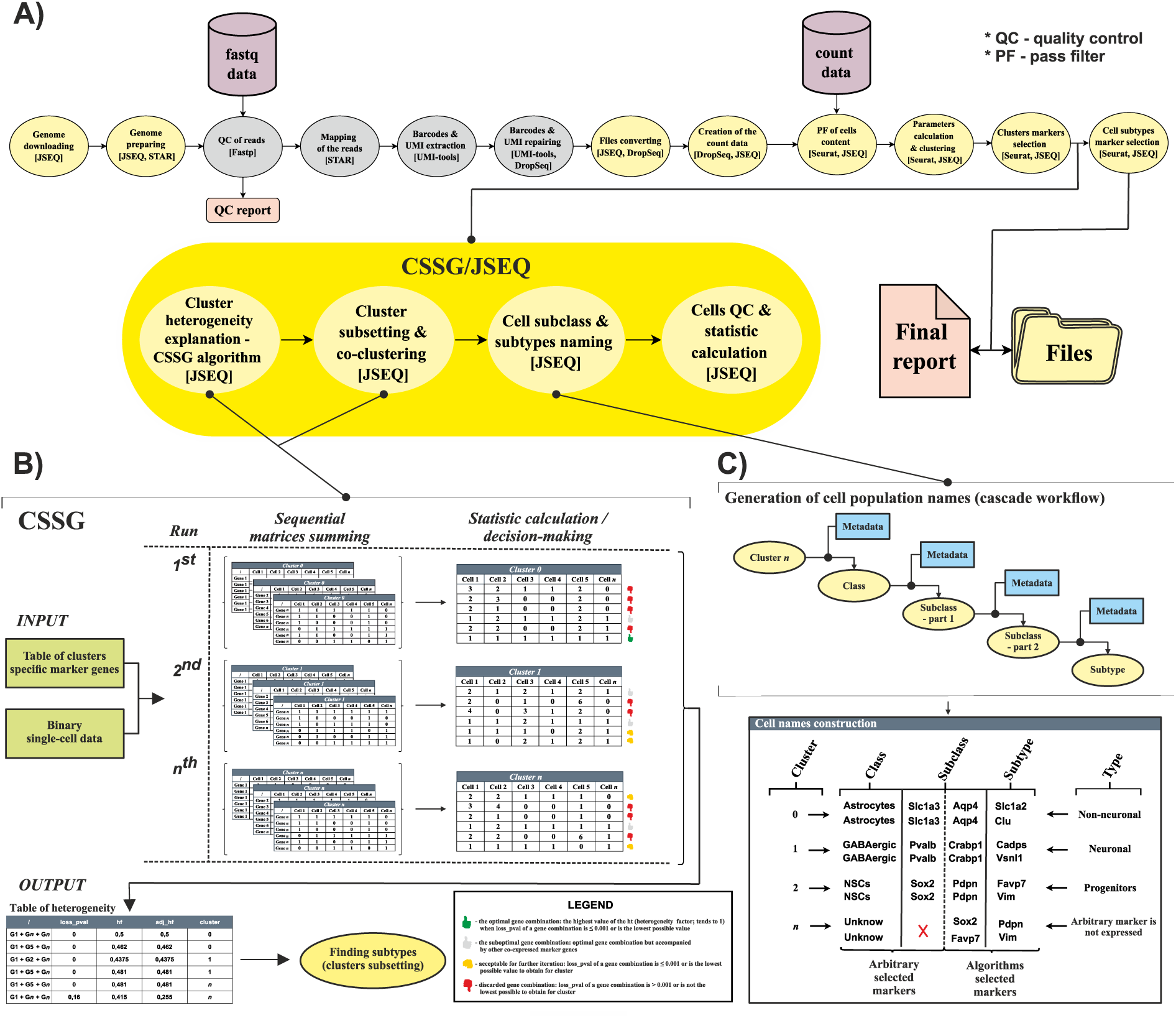
CSSG/JSEQ general workflow, CSSG algorithm details, and cell population naming strategy. A) Graphical scheme of CSSG/JSEQ pipeline workflow. Two major data entry points in the JSEQ pipeline, raw read data entry point in fastq format or pre-analyzed count data table entry point in the workflow; brief CSSG_®_ algorithm working scheme and implementation steps; B) scheme of the CSSG matrix-based iterative and algorithmic decisive approach and visualized means of statistics calculation to obtain the final results; C) Original method of cell population naming generated for high-resolution CSSG results. All names consist of descending detail degrees, and the final name is subdivided into class, subclass, and subtype parts. The mean of cell population naming is based on a cascade where consecutive name parts are selected based on metadata files.

### Pipeline phase: data preparation and generation of the count matrix

In case the entire pipeline is initiated from raw sequencing reads, the genome download module must be run first. In the JSEQ options, the user can select “human” or “mouse” or “custom” genome. JSEQ offers automatic download of the genomes from GENECODE, versions GRCh38.p13 for human or GRCm38.p6 for mouse [https://www.gencodegenes.org/] (8). The option “custom” offers additional possibilities, such as setting the download of other genomes versions or species (from GENECODE, NCBI, and ENSEMBLE). The “custom” option requires editing a simple text configuration file. The file also contains other options, such as switching on and off quality control of bases in sequencing results (in case of lack of quality information in fasta files) and the annotation enrichment by a specialized tool.

The analysis starts with quality control of the reads [fastp v.0.22.0] (9), which provides trimming of the adapters, bases quality control, minimal length control (important for UMI and barcode information included in Read1), and poly base tails trimming. Subsequently, genome indexing and mapping are performed by STAR v.2.5.0a, followed by double quality control. The control is an important step that involves repairing UMI and barcode sequences by UMI-tools (10) and Drop-Seq v.2.4.0 (11)]. Finally, the matrix of counts is generated in sparse format [Drop-Seq v.2.4.0]. All the external applications are controlled by JSEQ functions.

### Pipeline phase: processing count matrix

The analysis at this stage starts after creating the count matrix in the previous phase or can be started with previously prepared data in the format of the count matrix as a separate project in JSEQ. The advanced analysis uses features from Seurat v.3.1.5 package (12) as a framework where we incorporated a range of JSEQ approaches (data exchange and algorithms) to discover rare cell populations in high resolution together with their lineage and parent population.

In brief, the stages of the count matrix analysis in CSSG/JSEQ are: 1) multistage quality control; 2) data normalization and standardization; 3) data dimensionality reduction with principal components (PC) analysis; 4) automatic adjustment of PCs number; 5) multistage clustering; 6) selecting cluster-specific (subclasses) genes; 7) selection of gene combinations explaining cluster heterogeneity (CSSG algorithm); 8) cluster subsetting and co-clustering; 9) selecting subcluster-specific genes defining subtypes; 9) naming of cells (optional; helps to understand the heterogeneity of generated populations intuitively); 10) data visualization; 11) generating a complexity report with data explanation (graphs, tables, and description). In detail and for the current analysis, the quality control steps consisted of 1) removing cells with less than 500 genes per cell; 2) removing contaminations with double or multiple cells for one barcode; 3) removing cells with unknown origin (typically less than 10 per 1000 cells in a cluster). Data normalization was performed by log(CPM+1) formula individual for each cell. Variable genes across datasets were estimated using the equal frequency method by variance stabilizing transformation. Principal components (PC) were calculated on scaled data, and the number of PCs for further analysis was automatically adjusted to the correct number. The adjustment uses the variability check of the explained variance in consecutive PCs. Next, the estimated PCs were used for clustering with an algorithm such as the KNN (k-nearest neighbor) with Jaccard distance and SNN (shared nearest neighbor) graph with Louvain modularity optimization algorithms incorporated in Seurat. Marker genes specific to obtained cell clusters were calculated based on the MAST algorithm (13). The UMAPs plots, heatmaps, bar plots, and other scatterplots were used for data visualization. More information about the pipeline and available functions are provided in JSEQ^®^ single-cell pipeline manual deposited on the GitHub platform.

## CSSG validation methods

In the study, we also conducted validation of our CSSG algorithm (i) an investigation of gene expression between subtypes and additional clustering for refining the estimation of population trajectories, and (ii) an examination of specific diversity of expression between Alzheimer’s disease genes to establish disease relations between cell subtypes and the degree of their involvement in the disease. For comparing deregulated expression (DE) of genes between subtypes, we used the direct difference in expression [subtype 1^st^ normalized expression subtracted from subtype 2^nd^ normalized expression] and Wilcoxon-Test. We used hierarchical clustering with the ward.D algorithm to draw relations between particular subtypes for both analyses of the trajectories and the Alzheimer’s related genes. Analysis and diagrams were prepared in RStudio Version 1.3.1093 with R version 4.0.3.

## Results

### The preconditions for the CSSG algorithm

We developed a unique new scRNAseq CSSG^®^ algorithm (Cell Subtypes Selection by Genes) which is nested on our JSEQ^®^ platform and was prepared and customized for running this algorithm (combined name: CSSG/JSEQ). The algorithm and the pipeline provide a unique algorithmic decisive approach for preparing very high-resolution tissue cell composition in terms of populations and cell intermediate states. The pipeline is designed to maximize the amount of input scRNAseq data and applies various control steps to reach the maximum possible data quality. Moreover, the obtained data can be modified and/or used for further analysis with little effort. Finally, it meets the requirements of everyday researchers with little IT background for a scRNAseq tool that provides a high degree of automatic processing. In addition, our naming approach creates names for cell classes, subclasses, and subtypes, showing in the name of the population the cascade of names for the most important marker genes. The such naming principle can also demonstrate trajectories between intermediate cell states.

Our CSSG helps to obtain more efficient results from single-cell sequencing containing lower numbers of cells and/or lesser depth of data sequencing. To achieve this, we performed a series of tests with different datasets (different sequencing depths, different numbers of cells, different degrees of tissue development, etc.) and established a course of action to maximize the obtained results using additional analysis features. To confirm the superiority of our algorithm, we also conducted many validating analyses (described in the methodology).

### Decisive and automated CSSG^®^ algorithm for high-resolution scRNAseq analysis

The classical single-cell analysis is sufficient for many applications; however, in such a method, the obtained clusters still contain unexplained heterogeneity (cell subpopulations). The information is consequently hidden and can not be reliably discovered without specialized approaches. Moreover, we know the influence of many genes on specific cellular processes, the precise range and scale of gene expression levels, and the combinations that govern cellular processes are primarily unknown. Hence we do not fully understand the influence of subtle gene expression levels on rare population formation; we often discard information on subtle changes and discard cell populations that may be significant. Therefore we created CSSG^®^ algorithm (Fig. 1B), a very powerful algorithmic tool that discovers the diversity of individual gene occurrence inside initial cell clusters and allows for the generation of smaller, less heterogeneous subsets, which we call “subpopulation”. In the CSSG approach, we decided to shift our focus from information about the single gene expression levels to the presence of the collective sets of gene expressions in single cells. We transformed expression levels in each cell inside a particular cluster into binary data, where the value ‘1’ denotes the presence of the gene expression, and ‘0’ represents the lack of gene expression in a single cell belonging to one cluster.

Each iteration of the CSSG algorithm focuses on a single cluster, inside which starts the next sequence of iterations that explains the heterogeneity of the cluster. Starting the sequence of iterations inside the cluster requires a pair of data matrices. The first matrix simply consists of repetitions of one vector, which contains binary data for one gene for each cell in the cluster (# of cells is the # of columns). The second matrix contains binary data for each gene (row) and for each cell in the cluster (columns), excluding the data for the gene repeated in matrix 1. Then, the summing of two matrices (equation 1) is performed and repeated until all combinations of two genes in the cluster are obtained. Subsequently, the action of CSSG obtains all possible combinations for 2, 3, or more genes using binary matrices and calculates relevant statistics. When appropriate combinations of genes are established for all clusters, the algorithm returns a table of the best combinations explaining heterogeneity for each cluster, which are then used in cell subtyping and naming.

The intuitive principle of adding a par of matrices results in the matrix, where the co-occurrence of two or more genes in a single cell in the examined cluster is denoted as a multiplicity of “1” value (e.g., 2, 3, 4, *n*). For example, the presence of only one gene out of all genes in the current iteration in one cell is denoted as “1,” and the expression lack of any is denoted as “0”. The value 2 or greater present in the matrix (co-occurrence of two or more genes in one cell) does not explain the heterogeneity and is obviously not favorable for discovering rare populations; it may, however, constitute an intermediate form between cell subtypes. The possible number of two gene combinations (*C_n_*) in one cell is between 1 and the factorial 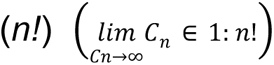 where *n* is the maximum number of marker genes inside the analyzed cluster. To improve the performance of the CSSG, it uses only preselected by MAST algorithm marker genes to each cluster at a significant level *p* <= 0.05 of the Wilcoxon-Test. It is one of the elements that greatly reduces computation time, CPU, and RAM usage.

The CSSG algorithm network in each iteration calculates the specific statistics that serve as a weight, limiting the amount of data in iterations because the number of combinations in the successive iterations tends to infinity. The following statistics parameters: *loss_pval* (equation 2) [assumpted value describing the number of cells discarded during cluster analysis]. According to the *loss_pval* parameter, cells are discarded when no marker gene occurs in these cells within the analyzed cluster. The user can also change the *loss_pval* parameter reflecting the principal weight for the CSSG algorithm network that the expected combination of genes (heterogeneity) for an analyzed cluster was reached, and that algorithm can stop processing the analyzed cluster (stop point) If the algorithm can not reach an equal or lower value than the user-defined *loss_pval* (default 0, 001); it will search a combination with the lowest *loss_pval* that can be obtained. For example, if in a cluster containing 100 cells and one cell can not be described by any marker genes, the minimal loss_pvale which can reach is 0.01, so the algorithm would iterate infinitely and never finish. Another important parameter governing the CSSG algorithm is the *ht* (equation 3) [heterogeneity factor] which is a value informing the algorithm about the level of heterogeneity of the cluster explained by the genes combinations. The ht is also one of the wights for algorithm network influencing further decisions of the algorithm during analysis which is explained in the subsequent paragraph. The *adj_ht* (equation 4) [adjusted heterogeneity factor] is the heterogeneity factor value adjusted by the value of discarded cells from the cluster. The aim of the *adj_ht* is to introduce the compromise between maximal possible heterogeneity and the number of discarded cells within the cluster.

In the process of finding the optimal heterogeneity, subsequent combinations of genes are forwarded to the next iteration of the analyses based on algorithm decision evoked by the *loss_pval* and ht parameters. Such parameters obtained for all gene combinations are used to select weights using quartile statistics (Q25 – first quartile and Q75 – third quartile). The first weight for the algorithm network is determined when all combinations of the two genes are achieved. At this point, only combinations of genes with ht value greater than the third quartile of all estimated ht values [*ht* >= Q75 of all ht] are subjected to further analysis steps. When including three and more gene combinations are required to explain optimal heterogeneity, the *loss_pval* determines the weight for the transfer of data to further analysis. The weight is calculated based on the first quartile of all *loss_pval* statistics [*loss_pval* <= Q25 of all *loss_pval*].

Used weights are necessary to reduce the number of results to only highly relevant combinations. Although the algorithm is optimized for multiprocessing, the excessive number of gene combinations can extend the algorithm’s working time and may lead to CPU and RAM overload. If all conditions are reached, and all sensitive gene combinations are determined in the analysis, the results are saved to the final result table of clusters heterogeneity with their statistics.

Based on the table of cluster heterogeneity, the cluster sub-setting is conducted inside each cluster using the dominant expression [max(log(CPM+1)] of genes from the best genes combination based on the *ht* or *adj_ht* statistic for a given cluster. The user can decide which parameter is best suitable for the current single-cell experiment depending on the required resolution level. The cell subtyping can be conducted according to both parameters, including all consequences they cause and are described above. The default setting is *adj_ht* to minimize cells discarded from the cluster. The minimum number of cells in the subtypes is checked using the binomial test at a significance level (0.1 in the case of very rare sub-populations). Cell subtypes with lower p-value are excluded from the analysis. All parameters for CSSG and for subtyping workflow can be set in the configuration file.

#### Matrices operations

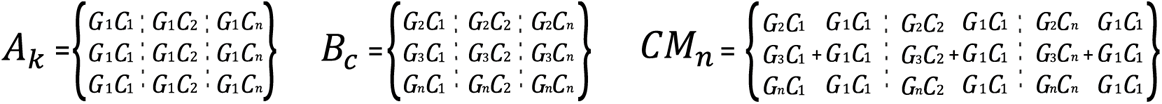

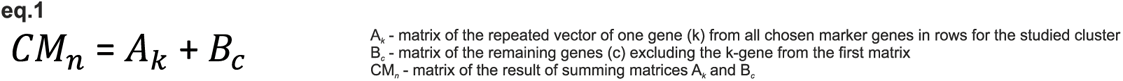

#### Vector of genes combination

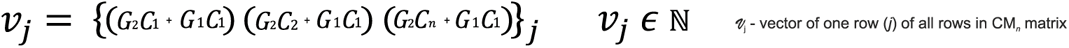

#### Equation conditions

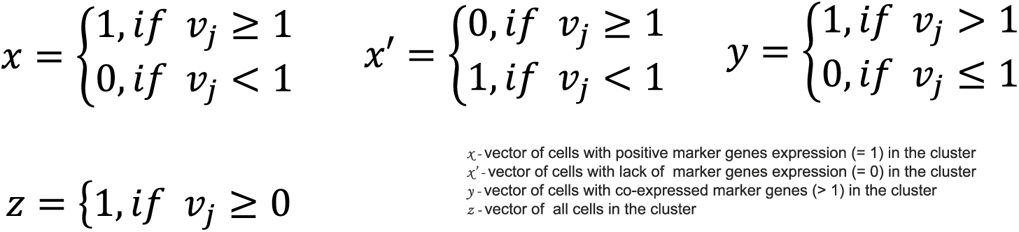

#### Equations

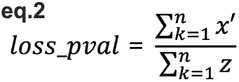

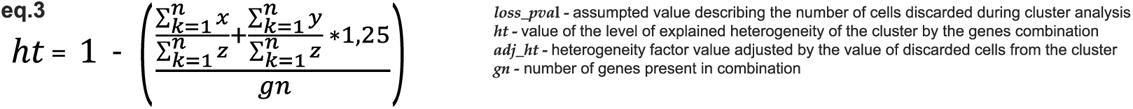

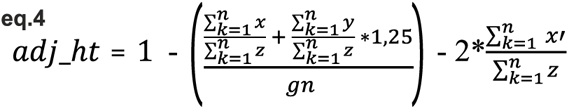

### Cell population naming algorithm

We developed a cell population naming algorithm to support our CSSG methods. The algorithm is responsible for establishing the naming of the received cell classes (class name), subclasses (two marker genes of the subclass), and subtypes (single marker gene of the subtype). The advanced algorithm has been designed to create systematics of the cell population complexity in analyzed tissue. The algorithm always requires two tables, the table containing marker genes metadata (differing for each step of naming; class, subclass, etc.) and the other table containing expression levels [mean(log(CPM + 1))] of all genes found in metadata in all clusters (for naming class and subclass) or all subtypes (for naming subtypes). In the process of cell naming, the algorithm selects the gene with the maximal expression for each cluster or each subtype, and subsequently, the algorithm screens metadata to name the cell population defined by this gene. In the case of naming class and the first part of subclass name (a gene name), the metadata table contains arbitrary predefined and most current state-of-the-art literature and databases of cell marker genes. For the second part of the subclass name, the cell naming algorithm uses the table of cluster-specific markers selected by the MAST algorithm. Finally, for naming subtypes, the algorithm uses genes from the table of heterogeneity obtained from the CSSG algorithm. Figure 1C summarizes the naming approach which is based on the cascade principle of gene tables and information in the metadata table, where the final cell population name is composed of the cell class name, two genes names for the subclass, and the gene name for the subtype.

In addition, the brain as a model tissue for our pipeline and algorithm is a particular challenge since it is very complex and contains a multitude of regions and cell types. Therefore especially for the brain tissue, we constructed an enhanced metadata table with markers for different brain subclasses belonging to neuronal, non-neuronal, and progenitor cell types. The neuronal part of cell names is based on genes involved in particular categories. The categories are: signaling related to neurotransmitters (Tab. 1), signaling by neuropeptides (ligand or receptor) or calcium-binding proteins (Tab. 2), and by neurotransmitter released such as: excitatory (glutamate), inhibitory (GABA, glycine), and modulatory (choline, noradrenaline, serotonin, dopamine, and nitric oxide). In addition, for the cerebellum, we can distinguish Purkinje (inhibitory), Golgi (inhibitory), and granule (excitatory) cell classes. In the case of non-neuronal and progenitor cell names, the first subclass marker used is the one that defines the cell type from table 3 (e.g., Oligodendrocytes Olig2, NPCs Sox2, etc.).

**Table 1.**
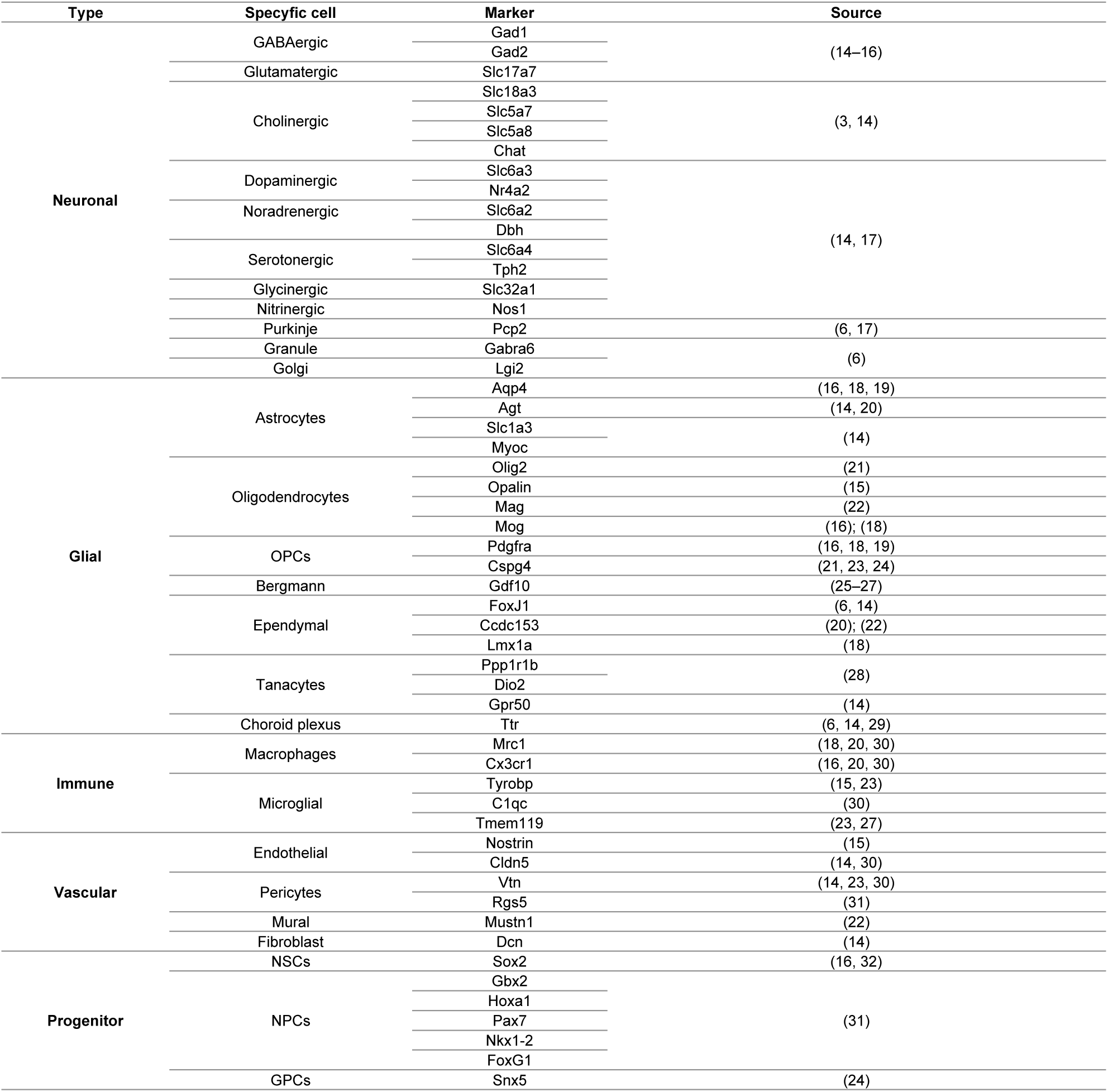
Table of genes that are markers for brain cell populations (neuronal and non-neuronal) selected by extensive state-of-the-art literature-mining. The marker genes of the cell populations were selected according to literature reports assuming that the marker gene is unique through the brain. Brain region-specific markers were avoided. The table was used as the metadata for generation cell class or cell class/subclass name parts.

**Table 2.**
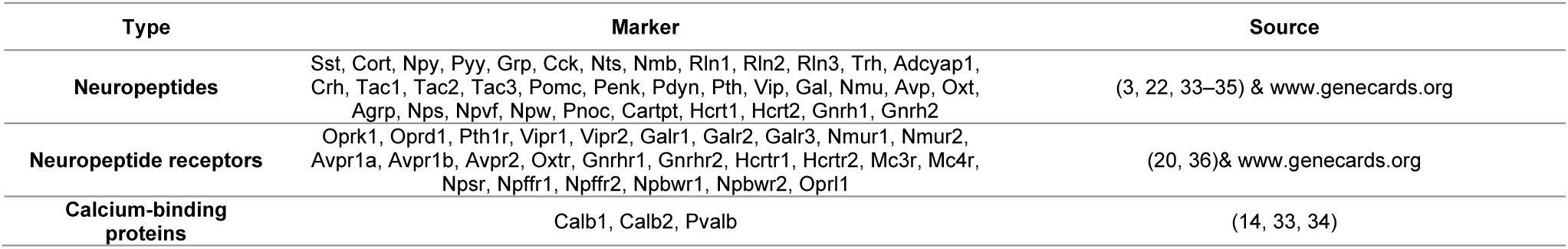
Table of genes that are exclusively markers for neuronal brain cell populations selected by extensive state-of-the-art literature-mining. As in Table 1, the marker genes of the cell populations were selected to be unique through the brain. The table was used as the additional metadata mainly for generation cell subclass name parts of neuronal cells.

**Table 3.**
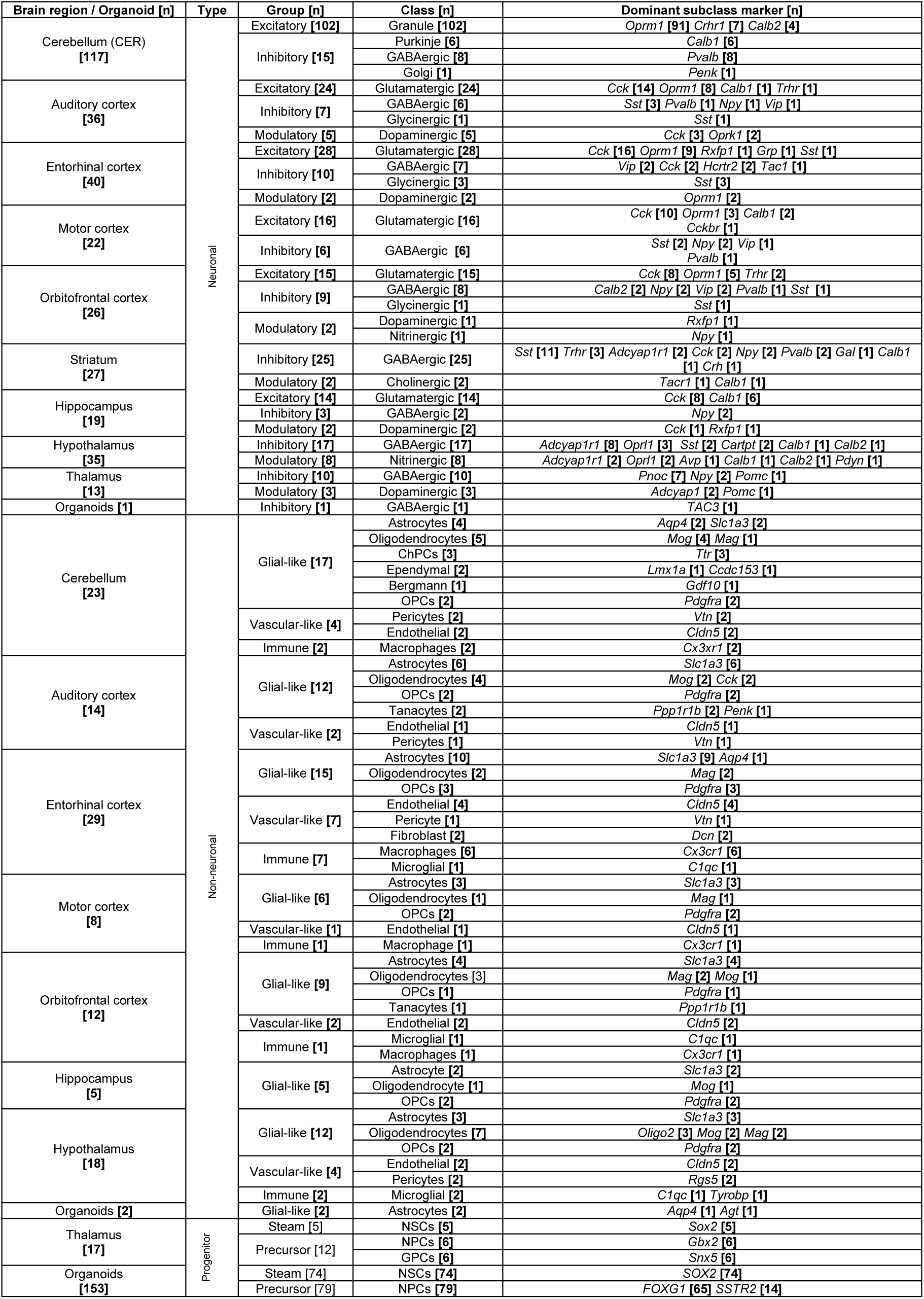
The summary of high-resolution cellular brain complexity obtained by CSSG/JSEQ. The table contains neuronal, non-neuronal, and progenitor cell populations of all analyzed brain regions and their dominant markers for subclass. The numbers in brackets indicate numbers of cell subtypes (lowest level of cell population in cell cluster) attributed to brain region, class, and arbitrary groups.

### CSSG validation by examination of the global complexity of the brain and human brain organoids

We validated our CSSG algorithm results by conducting an extensive analysis of the mouse brain and human brain organoids. In the case of brain structures, the algorithm discovered many to-date unrevealed cell population subtypes. To examine the validity of brain complexity, we plotted the relations of the number of cells, the number of subclasses, and the number of subtypes (Fig. 2). We always found linear relations in the case of fairly mature populations. The relations of the cells vs. subclasses are also linear for organoids, similar to mature cells. Interestingly there is no more linearity for organoid cells when cell vs. subtypes or subtypes vs. subclasses are plotted. Such behavior confirms that in the case of human brain organoids, most cells are in the stage of intensive development, which results in increased heterogeneity of data and a greater number of subtypes obtained. On average, we have detected 24 subclasses which created on average of 62 subtypes with the exception of the human brain organoids, where we detected 22 subclasses and 157 subtypes. Together, this demonstrates our algorithms’ advancement in explaining cellular tissue heterogeneity.

**Figure 2.**
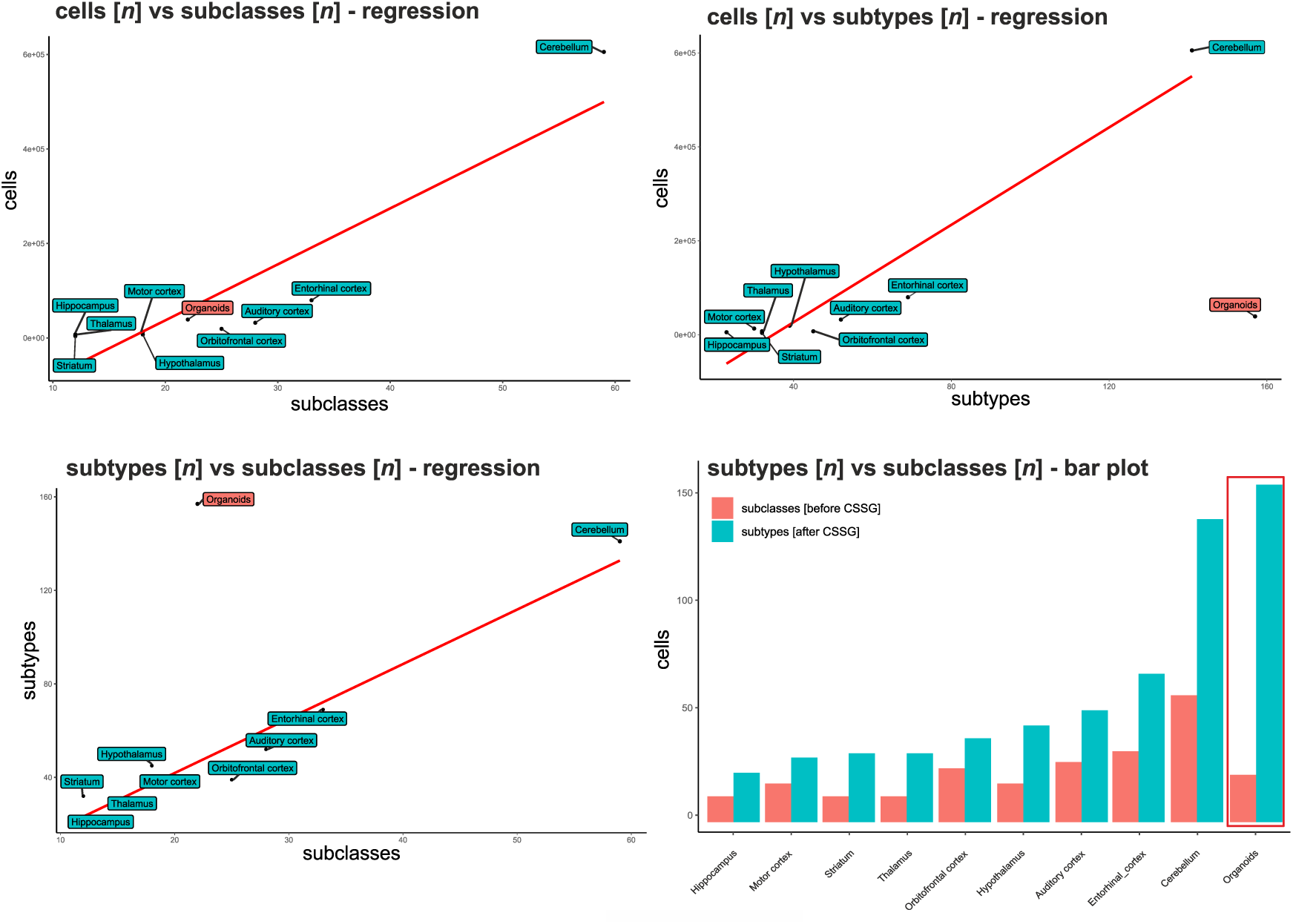
The diagrams demonstrate the relationship between the numbers of cells, subclasses, and subtypes before and after using the CSSG algorithm. The number of cells vs subclasses diagram (before using the CSSG algorithm) reveals the linear relation for all datasets. The CSSG analysis causes a dramatic enrichment of results by introducing new populations called “subtypes”. This process increases cellular resolution and knowledge about the cellular structure of the tissue. The subtypes are characterized by lower heterogeneity of cells belonging to a single subtype; therefore, we obtain high consistency of cellular composition. In addition, the analysis after the CSSG approach (cell number vs a number of subtypes and subtypes vs cell classes) reveals an extreme example of an organoid where there is no linearity with brain regions and where CSSG discovered very many subtypes. This indicates that particularly immature tissues, such as organoids, exhibit particular diversity.

Overall, we obtained 239 cell subclasses and 620 subtypes (Tab. 3) in the analysis. In neuronal cell types were 326 subtypes, including 200 excitatory, 102 inhibitory, and 24 modulatory subtypes. The non-neuronal cells contained 124 subtypes consisting of glial-like (astrocytes and oligodendrocytes etc.), vascular-like, and immune cells. We also identified many progenitor cell subtypes (170) prominent in the thalamus and human brain organoids.

Interestingly, the striatum, hypothalamus, and thalamus contain a variety of cell subpopulations characterized by markers that define neurons with double excitatory and inhibitory roles such as dopaminergic, cholinergic etc.

In all neuronal types, we identified many subpopulations characterized by important neuropeptides, their receptors, and calcium-binding protein such as Sst (24 subtypes), Pvalb (13), Npy (11), and Adcyap1r1 (10), Oprm1 (116), Cck (56) and Calb1 (9) Cck (4), Adcyap1 (2), Adcyap1r1 (2), Calb1 (2), Oprk1 (2), Oprl1 (2), Oprm1 (2) and Rxfp1 (2).

Together, we demonstrated differences between obtained cell subtypes, greatly increasing the resolution of populations and improving brain complexity knowledge. Examples of clearly improved cluster separations are shown in Figure 3. The entire subclass pattern and the resulting subtypes can be checked in supplement 1.1. Detailed tables describing the composition of the subtypes with markers and extended enrichment conducted for genes presented in the subtypes in supplement 1.2.

**Figure 3.**
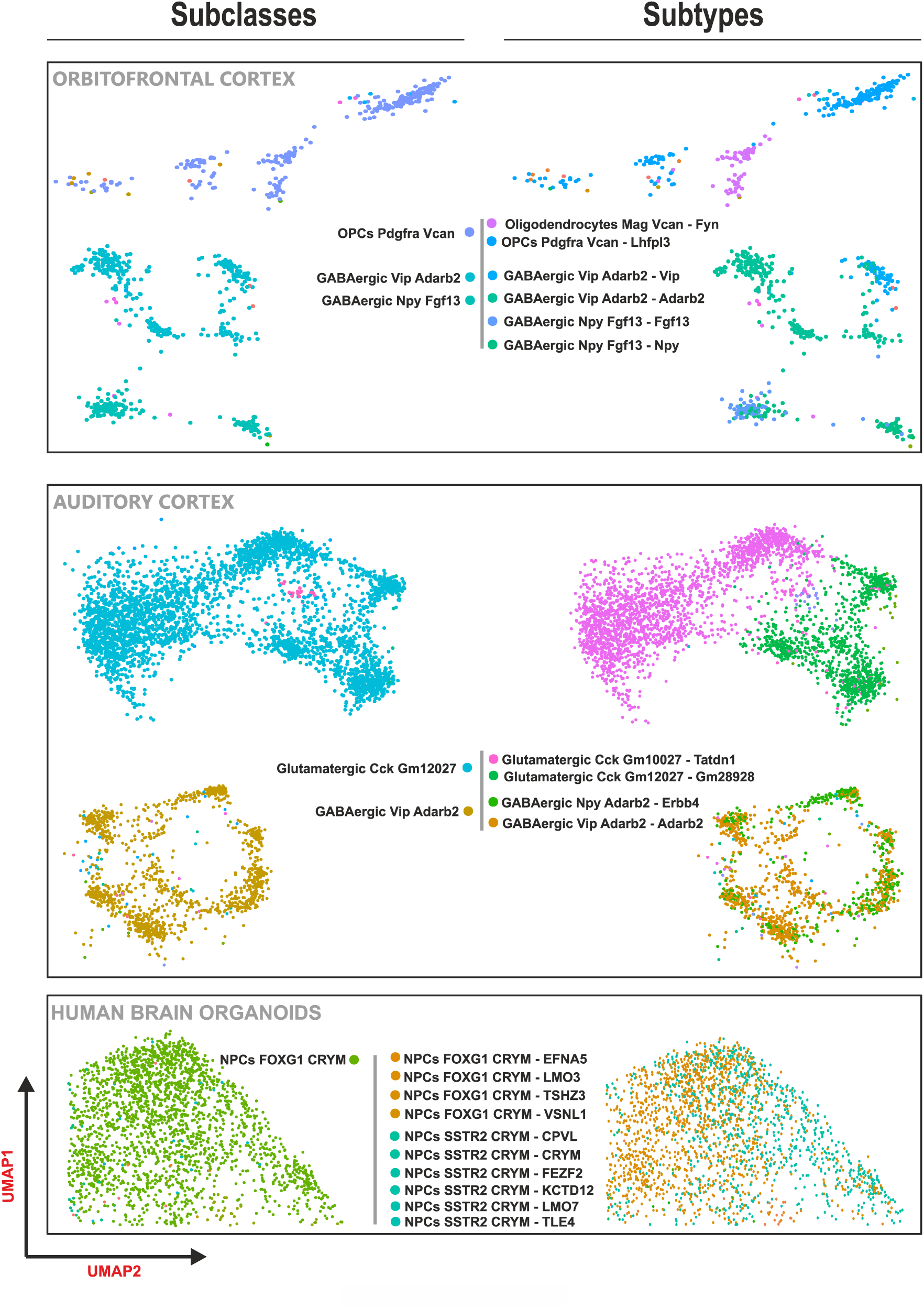
The CSSG/JSEQ algorithm has the ability to perform the one-step-separation of an initial cluster comprising only subclasses (used in traditional low-resolution analysis) to multiple cell subtypes and their genetic markers from the brain (OCTX, ECTX) scRNAseq and human brain organoids. Note that a single cluster can produce unbiased diversity by CSSG only by decisive action. The data are visualized using parts of the UMAP plots.

### CSSG/JSEQ in cell fate and trajectory discovery

Most importantly, the CSSG/JSEQ performs exact mining of genetic markers in the initial cell clusters which initially seem to have a uniform differentiation status. The resulting subtypes isolated from the initial cluster have a combination of markers that allows separating into specific populations with substantially distinct differentiation and maturation stages. For instance, the subtypes showing the same second marker of subclass in cell name (Fig. 4) and belonging to one initial cluster are now assigned to different classes, namely oligodendrocytes and OPCs. This demonstrates the high-resolution capabilities of the pipeline to generate the trajectories of cells, such as oligodendrocytes showing detailed cell fate. In addition, we used all identified oligodendrocyte populations from the cerebellum, hypothalamus, orbitofrontal cortex, and hippocampus and conducted hierarchical clustering of their subtypes. Moreover, based on literature collection including single-cell experiments solely based on the analysis of oligodendrocytic fates, we established precise developmental and differentiation status of individual oligodendrocyte subtypes named OPC (oligodendrocyte precursor cells), COP (differentiation-committed oligodendrocyte precursors), preMO (pre-myelinating oligodendrocytes), NFO (newly formed oligodendrocytes), MFO (myelin-forming oligodendrocytes), and MOL (mature oligodendrocytes) in previous works. We confronted our subclass and subtypes and their precise markers obtained by CSSG/JSEQ with the collection of previous works. We noticed that our subclass and subtype markers’ information, arranged as a cascade of gene names (Fig. 1C), reflecting the intermediate oligodendrocyte cell maturation stages, visually demonstrated as the hierarchical tree (Fig. 4).

**Figure 4.**
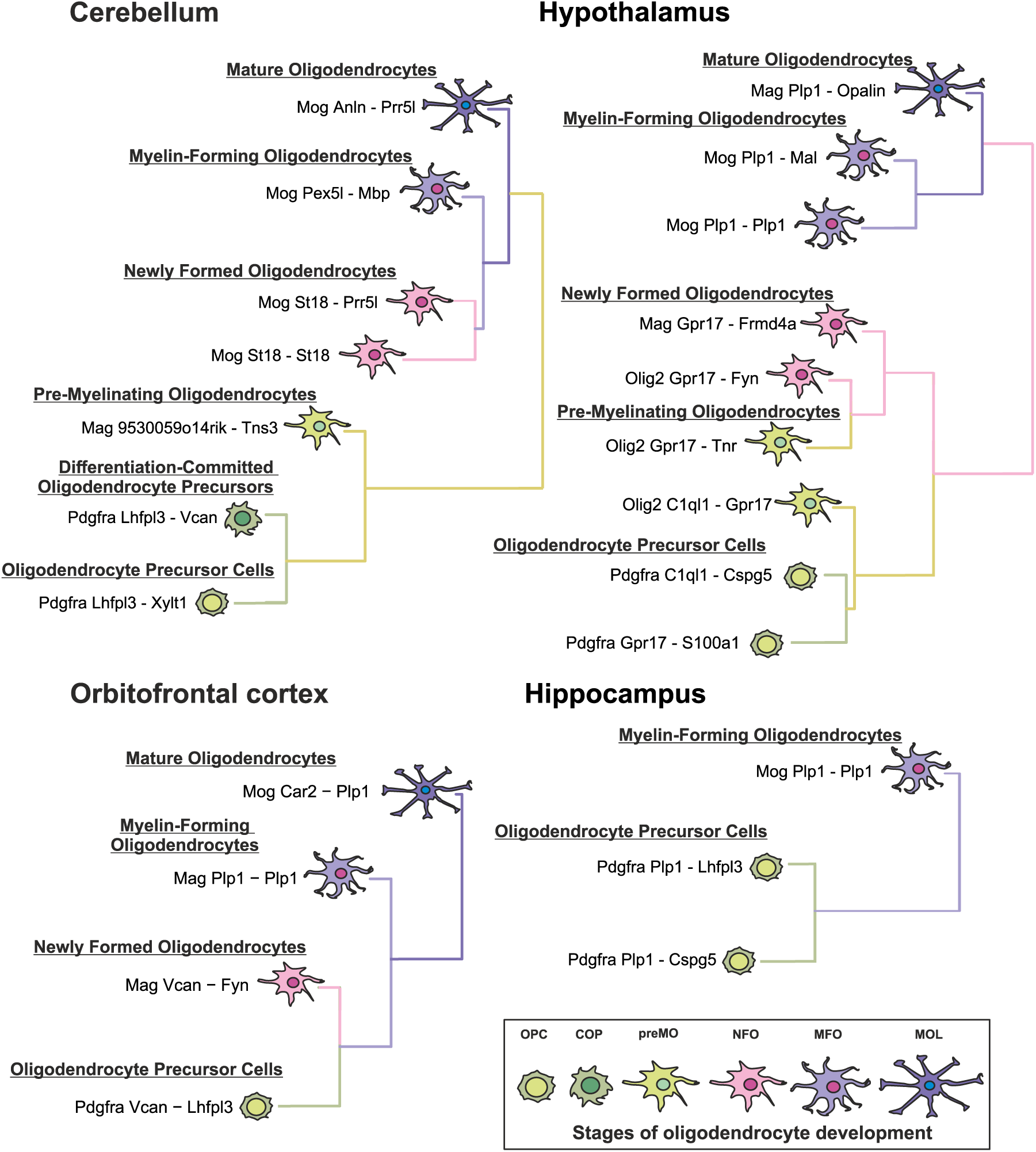
The CSSG/JSEQ algorithms are suitable for demonstrating unbiased trajectories of cell development. The figure demonstrates that CSSG analysis data suggested oligodendrocyte subtypes as their intermediate stages, revealing the trajectory of oligodendrocyte development. The identified cell types and their markers were confronted with state-of-the-art literature data. The CSSG-generated data confirmed that identified populations were not described to date or were described before but were not linked by trajectory previously, or their relations were already reported.

### CSSG/JSEQ algorithm in AD-related genes analysis

We also used the CSSG/JSEQ outputs to examine the suitability of our approach to biological problems focused on important genes involved in Alzheimer’s disease. We analyzed the distribution of Apoe, App, Mapt, Psen1, Psen2, and Trem2 genes in the cell population globally within brain regions: hypothalamus [HYP], thalamus [TH], motor cortex [MCTX], auditory cortex [ACTX], orbitofrontal cortex [OCTX], entorhinal cortex [ECTX], hippocampus [HIP], cerebellum [CER], and human brain organoids [HBO] (supplement 1.3.1) and then focused on differences in AD gene expression among the subtypes belonging to one initial cluster (supplement 1.3.2).

Analysis of the gene distribution across the different mouse brain regions and HBO shows that Apoe is most expressed and strongly related to Astrocytes and OPCs in the OCTX, MCTX, ACTX, ECTX, TH, HYP, STR, CER, HBO (Fig. 5, 6, 7); oligodendrocytes in the MCTX, OCTX, ECTX, ACTX, HIP, HYP (Fig. 5, 6) and ependymal cells in the CER (Fig. 7). Interestingly most data sets coherently demonstrate that high expression of Apoe is present in non-glial cells and neurons, such as in vascular cells in the MCTX, OCTX, ECTX, ACTX, HIP, STR, HYP, CER (Fig. 5, 6, 7); immune cells in the OCTX, ECTX, HYP, CER (Fig. 5, 6, 7), and glutamatergic neurons in the OCTX, ECTX, ACTX, HIP (Fig. 5). Finally, Apoe expression is also present in precursors, namely in GPCs (TH, HBO) (Fig. 6, 7) and NPCs in the HBO (Fig. 7).

**Figure 5.**
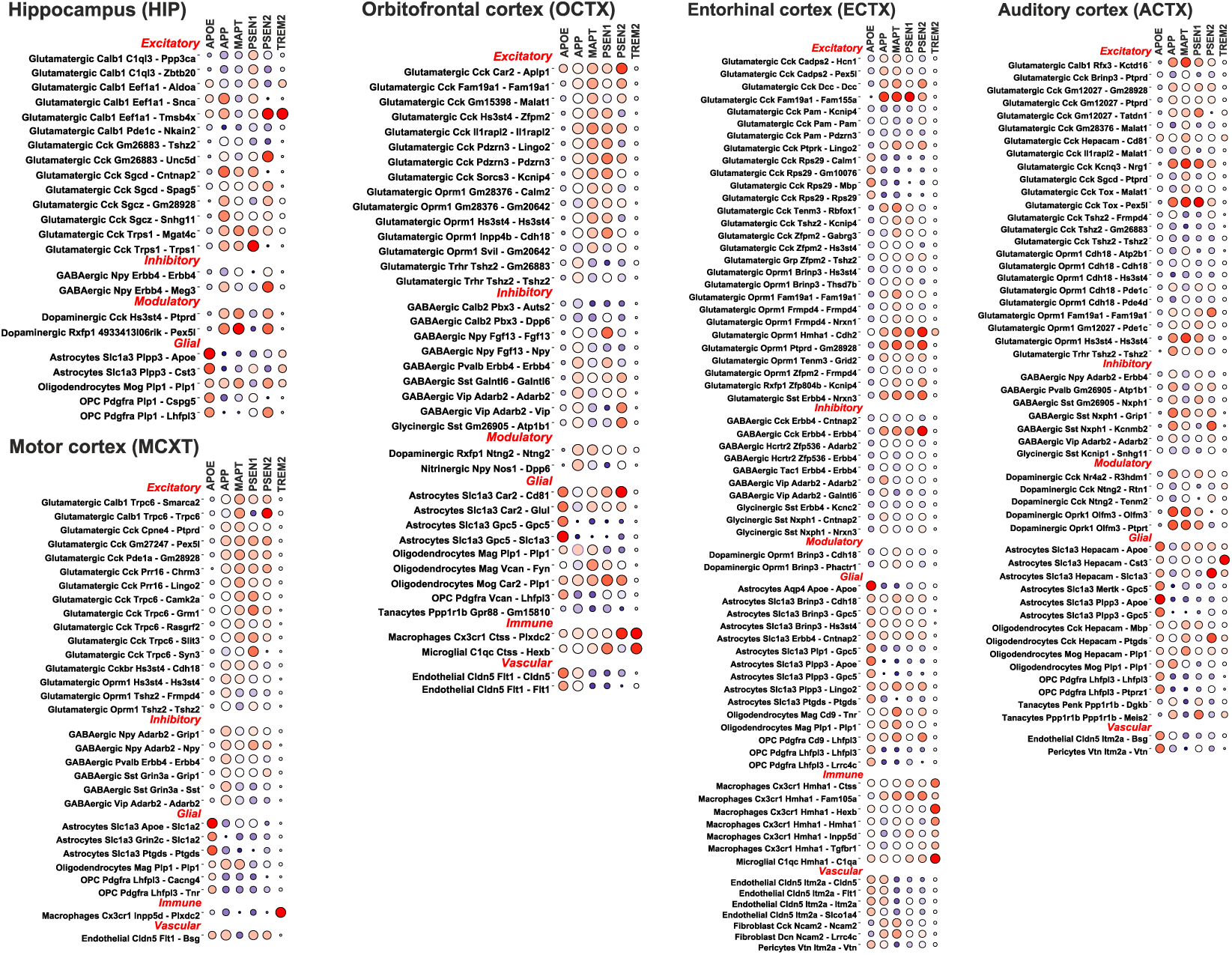
The CSSG/JSEQ, in one step, determines high-resolution brain cell subtypes from hippocampus and cortex areas selectively containing gene transcripts with an inevitable role in AD development. The selection of the canonical AD gene allows to mine of the global diversity of cell populations that can be affected by AD in the hippocampus (HIP), motor cortex (MCTX), orbitofrontal cortex (OCTX), entorhinal cortex (ECTX) and auditory cortex (ACTX). Note that CSSG determined many neuronal populations where the occurrence of transcripts for App, Mapt, Psen1, and Psen2 is highly correlated. Increased expression of the Apoe gene is characteristic of populations of astrocytes, microglial endothelial, and pericyte cells. Trem2 is present in mouse brains in microglial populations and sometimes correlates with the Apoe gene. The CSSG also identifies populations with the composition of AD genes that was not described before.

**Figure 6.**
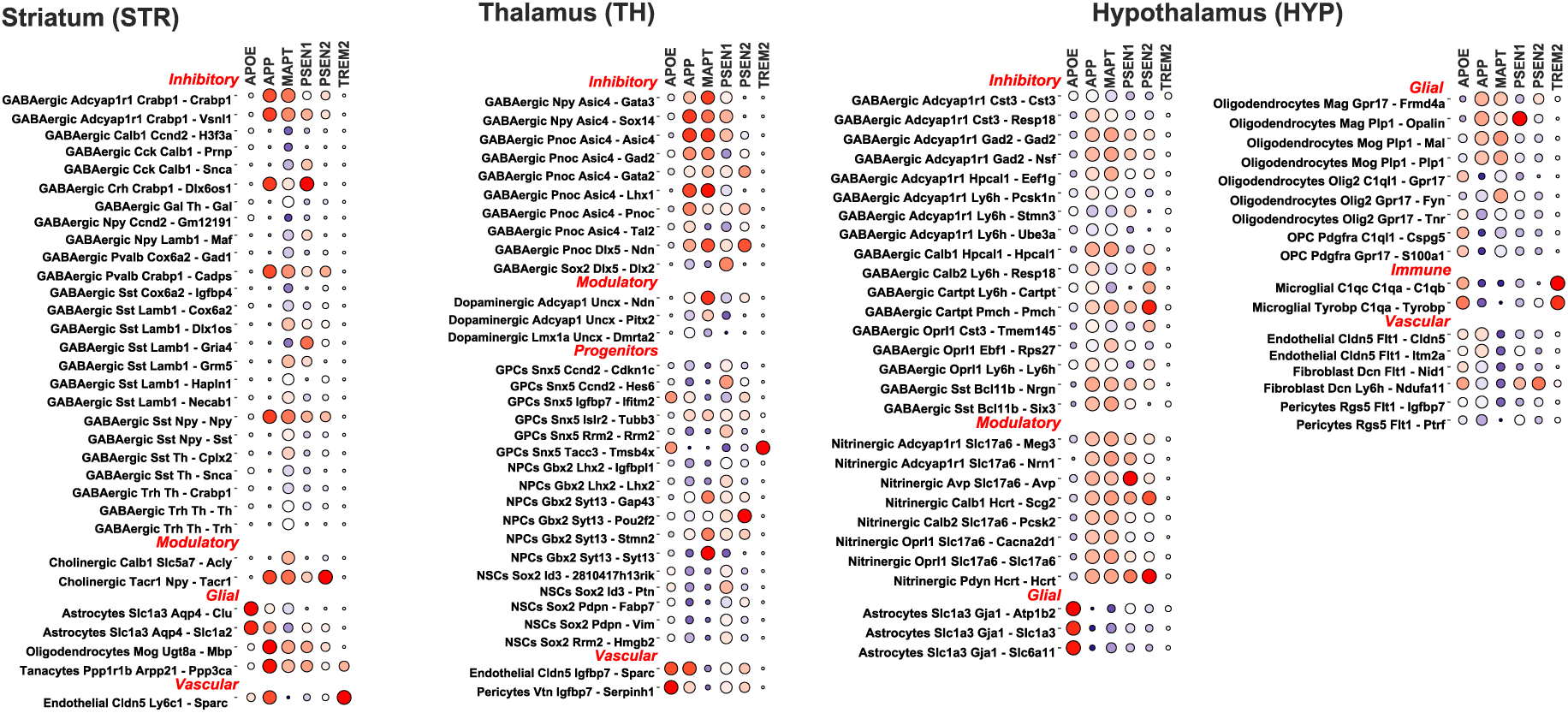
The CSSG/JSEQ, in one step, determines high-resolution brain cell subtypes from the striatum (STR), thalamus (TH), and hypothalamus (HYP) areas which selectively contain gene transcripts with an inevitable role in AD development. The brain regions contain many GABAergic cells which express AD genes. Again high correlation of App, Mapt, Psen1, and Psen2 occurs.

**Figure 7.**
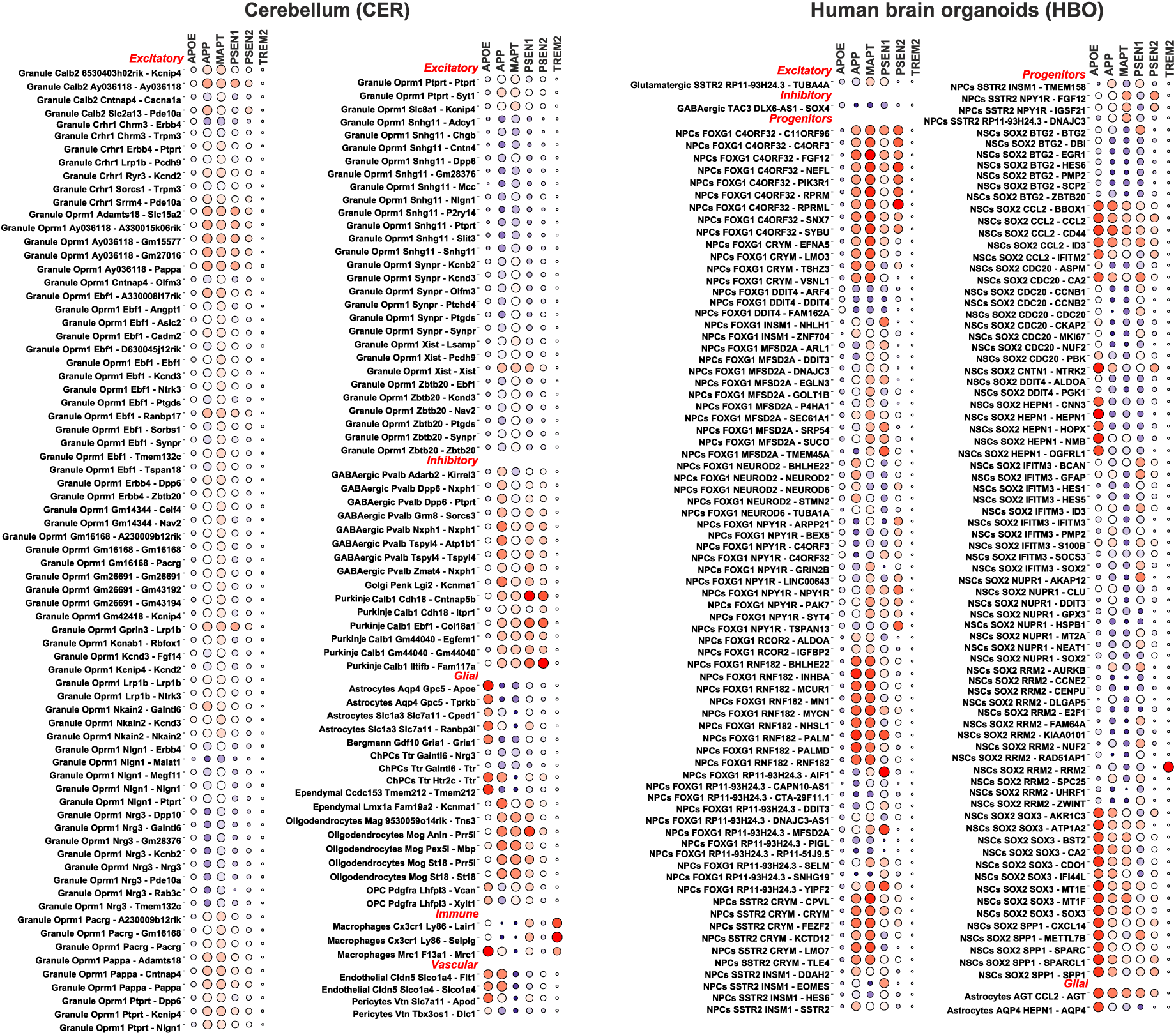
The CSSG/JSEQ, in one step, determines high-resolution brain cell subtypes from cerebellum (CER) and human brain organoids (HBO) containing gene transcripts with a role in AD development. Both scRNAseq datasets contain many cell subtypes. The cerebellum contains a large variety of subtypes; however, App, Mapt, Psen1, and Psen2 gene transcripts highly occur in inhibitory neurons such as Purkinje cells, while the occurrence in granule cells is lower. In the case of human brain organoids, cell populations mainly demonstrate an early developmental stage and divide between progenitors subtypes and subtypes with neuronal stem cell characteristics. Compared to subtypes of NSC, progenitors are already committed populations with APP, MAPT, PSEN1, and PSEN2 related to AD genes but do not coexpress APOE. The NSC subtypes in organoids often coexpress APOE together with APP, MAPT, PSEN1, and PSEN2.

Apoe is mostly not associated with Mapt, Psen1, Psen2, and Trem2 genes in cell populations. The partner of Apoe occurring the most is the App gene, which is visible in some subtypes of astrocytes, OPCs, oligodendrocytes, pericytes, choroid plexus, and glutamatergic cells in the whole brain (Fig. 5, 6, 7). The strong relation between Apoe and App expression is seen in one subtype of astrocytes, widely present in NPCs derived from HBO (Fig. 7) and also the strong partner of some endothelial cells. The connection of Apoe expression with other genes such as Mapt, Psen1, Psen2, and Trem2 is sparsely present, but it largely depends on the cell’s subclass and subtype.

The App gene is expressed with high frequency in neurons across the mouse brain (Fig. 5, 6, 7) and HBO (Fig. 7) and is highly accompanied by the expression of Mapt, Psen1, and Psen2. The combined correlation among Mapt, Psen1, and Psen2 expression highly depend on the cell subclass and subtype.

The increased Trem2 gene expression is very strongly related to immune cells and some astrocyte subtypes. In a few cases, we note an increased expression of Trem2 in glutamatergic subtypes in the ECTX, ACTX, HIP (Fig. 5), and GPCs in the TH (Fig. 6), NPCs in the HBO (Fig. 7), tanycytes in the STR, HYP, endothelial in the STR (Fig. 6), and oligodendrocytes in the ACTX (Fig. 5).

Thanks to the resolution offered by CSSG/JSEQ, we were able to conduct statistical and differential expression analysis of AD-related genes between cell subtypes within an initial cluster (subclass) (supplement 1.3.2), and we identified clusters containing subtypes that are the most heterogenous in relations to expression and a number of AD genes (Tab. 4). Strikingly, inside the most heterogenous clusters there are subtypes with a different class affiliation for instance subtypes occurring in OCTX, ECTX, STR and CER (Fig. 8 and Tab. 4). The change of affiliation may result from a few reasons. These may be related to i) processes that change cell lineages during development, such as various populations of oligodendrocytes and OPCs (Fig. 4, 8), ii) cellular responses resulting from endocrine or paracrine stimuli resulting from the interplay between several subtypes within a cluster, e.g., such as inflammatory responses potentially taking place in astrocytes and macrophages (Fig. 8 and Tab. 4); iii) incorrect assignment of subpopulations to the cluster as a consequence of clustering algorithms itself, e.g., Purkinje occurring among granule subtypes (Fig. 8 and Tab. 4). In all these cases, CSSG subtyping is the superior method to better understand cell interactions as well as improve data quality by picking up misallocated cells. We also carried out subtypes hierarchical clustering, which revealed dependences and their relations. Exact AD subtypes relations are presented in the discussion. The possibility of selective analysis of genes in exactly defined subpopulations in high resolution can contribute to marker discovery and a better understanding of AD disease pathogenesis and the interaction between cells.

**Figure 8.**
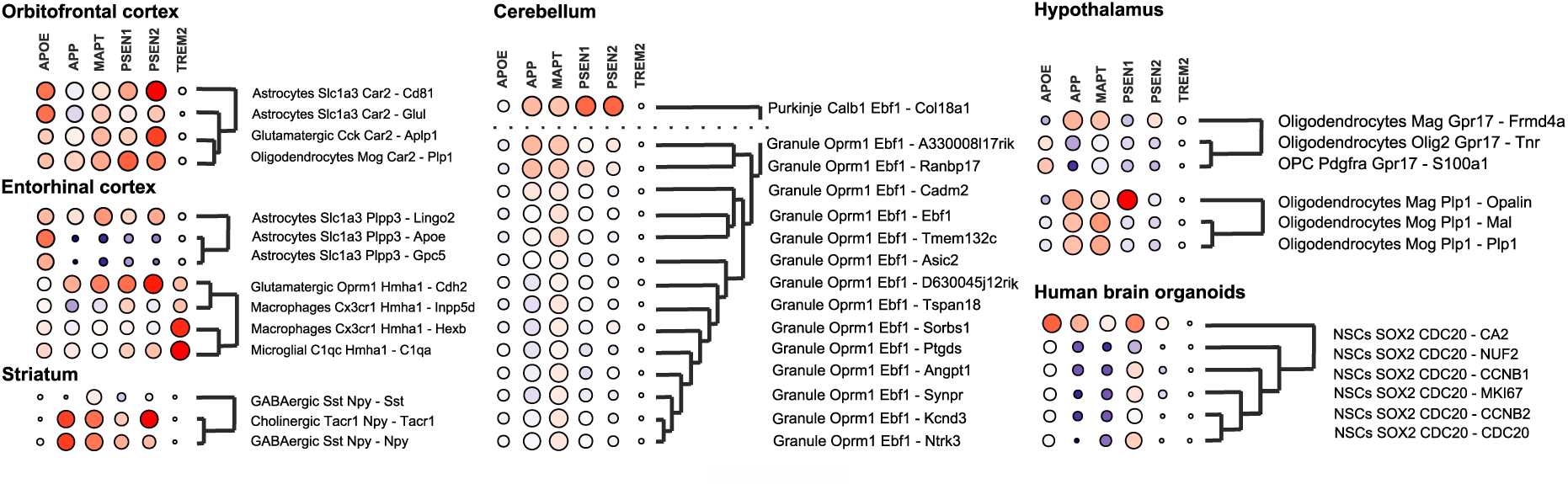
Relationships of populations of cell subtypes expressing characteristic genes related to AD, obtained by CSSG/JSEQ. Importantly the selected globally high-resolution subtypes (e.g., from OCTX, ECTX, STR, CER, HYP, and HBO) allow for tracking the subpopulation relations to other subtypes belonging to one initial cluster using hierarchical clustering. Inside the most heterogenous clusters, there are subtypes with different class affiliations, for instance, subtypes occurring in OCTX, ECTX, STR, and CER. The change of affiliation may be related to i) processes that change cell lineages during development (also presented in Figure 4), ii) cellular responses resulting from endocrine or paracrine stimuli resulting from the interplay between several subtypes within a cluster, e.g., such as inflammatory responses potentially taking place in astrocytes and macrophages (also in Tab. 4); iii) incorrect assignment of subpopulations to the cluster as a consequence of clustering algorithms itself, e.g., Purkinje occurring among granule subtypes (also listed in Tab. 4).

**Table 4.** Tabular representation of cell populations which show the most statistically significant differences in expression of AD genes (Apoe, App, Mapt, Psen1, Psen2, Trem2), while the expression differences in the single cell population must occur combined at the global level (region) and at the local level (cluster). The table contains subtypes belonging to different regions (OCTX, ECTX, STR, CER, HYP, and HBO). Such comparison of high-resolution subtypes allowed for identifying clusters containing subtypes that are the most heterogenous in relation to expression and a number of AD genes. These subtypes are compared to other subtypes within one initial cluster (local) using the Wilcoxon-Test statistics and real deregulated expression [numbers relate to subtype’s normalized expression, which is a difference of subtype 1st expression minus subtype 2nd expression]. The level of significance: [p-value: .> 0.05, * <= 0.05, ** <= 0.01, *** <= 0.001]. All results for other subtypes and regions are collected in the supplement 1.3.2 table.

| Region | Subtypes interaction | Subclass | ApoE | App | Mapt | Psen1 | Psen2 | Trem2 |
| --- | --- | --- | --- | --- | --- | --- | --- | --- |
| Cerebellum | Purkinje Calb1 Ebf1 - Col18a1 -> Granule Oprm1 Ebf1 - A330008\|17rik | Ebf1 | 0.21 . | -0.04 *** | 0.05 *** | 1.06 *** | 0.97 *** | 0 . |
|  | Purkinje Calb1 Ebf1 - Col18a1 -> Granule Oprm1 Ebf1 - Ranbp17 |  | 0.36 ** | 0.01 *** | 0.11 *** | 0.78 *** | 0.98 *** | 0 . |
|  | Purkinje Calb1 Ebf1 - Col18a1 -> Granule Oprm1 Ebf1 - Ebf1 |  | 0.14 *** | 1.03 *** | 0.66 *** | 1.14 *** | 1.18 *** | 0.003 * |
|  | Purkinje Calb1 Ebf1 - Col18a1 -> Granule Oprm1 Ebf1 - Tmem132c |  | 0.28 *** | 0.91 *** | 0.47 *** | 1.11 *** | 1.26 *** | 0 . |
|  | Purkinje Calb1 Ebf1 - Col18a1 -> Granule Oprm1 Ebf1 - Synpr |  | -0.02 * | 1.47 *** | 0.91 *** | 1.26 *** | 1.31 *** | 0 . |
|  | Purkinje Calb1 Ebf1 - Col18a1 -> Granule Oprm1 Ebf1 - Asic2 |  | 0.06 . | 1.04 *** | 1.02 ** | 1.23 *** | 1.27 *** | 0 . |
|  | Purkinje Calb1 Ebf1 - Col18a1 -> Granule Oprm1 Ebf1 - Ntrk3 |  | 0.003 *** | 1.15 *** | 0.81 *** | 1.18 *** | 1.25 *** | 0 . |
|  | Purkinje Calb1 Ebf1 - Col18a1 -> Granule Oprm1 Ebf1 - Kcnd3 |  | 0.03 ** | 1.2 *** | 0.81 *** | 1.17 *** | 1.3 *** | 0 . |
|  | Purkinje Calb1 Ebf1 - Col18a1 -> Granule Oprm1 Ebf1 - D630045j12rik |  | -0.02 . | 1.36 *** | 1.06 *** | 1.18 *** | 1.27 *** | -0.02 . |
|  | Purkinje Calb1 Ebf1 - Col18a1 -> Granule Oprm1 Ebf1 - Tspan18 |  | 0.21 * | 1.44 *** | 0.8 *** | 1.16 *** | 1.28 *** | 0 . |
|  | Purkinje Calb1 Ebf1 - Col18a1 -> Granule Oprm1 Ebf1 - Angpt1 |  | -0.09 . | 1.24 *** | 0.89 *** | 1.34 *** | 1.15 *** | 0 . |
|  | Purkinje Calb1 Ebf1 - Col18a1 -> Granule Oprm1 Ebf1 - Ptgsd |  | -0.39 . | 1.38 *** | 1.1 ** | 1.32 *** | 1.33 *** | 0 . |
|  | Purkinje Calb1 Ebf1 - Col18a1 -> Granule Oprm1 Ebf1 - Sorbs1 |  | 0.1 . | 1.42 ** | 0.93 *** | 1.22 *** | 1.03 *** | -0.02 . |
|  | Purkinje Calb1 Ebf1 - Col18a1 -> Granule Oprm1 Ebf1 - Cadm2 |  | -0.05 . | 0.68 *** | 0.78 *** | 1.15 *** | 1.26 *** | 0 . |
|  | Granule Oprm1 Xist - Xist -> Granule Oprm1 Xist - Lsamp | Xist | -0.17 ** | 1.12 ** | 0.86 . | 0.63 *** | 0.22 *** | 0 . |
|  | Granule Oprm1 Xist - Xist -> Granule Oprm1 Xist - Pcdh9 |  | -0.53 *** | 1.16 . | 1.31 *** | 0.68 *** | 0.26 *** | 0 . |
| Entorhinal cortex | Astrocytes Slc1a3 Plpp3 - Gpc5 -> Astrocytes Slc1a3 Plpp3 - Lingo2 | Plpp3 | 0.4 *** | -3.03 *** | -2.88 *** | -0.52 *** | -0.52 *** | -0.03 . |
|  | Astrocytes Slc1a3 Plpp3 - ApoE -> Astrocytes Slc1a3 Plpp3 - Lingo2 |  | 1.92 *** | -2.95 *** | -2.81 *** | -0.52 *** | -0.47 *** | -0.04 ** |
|  | Macrophages Cx3cr1 Hmha1 - Inpp5d -> Microglial C1qc Hmha1 - C1qa | Hmha1 | -0.92 *** | -1.29 *** | -0.29 . | 0.01 . | -0.21 . | -1.2 *** |
|  | Macrophages Cx3cr1 Hmha1 - Hexb -> Glutamatergic Oprm1 Hmha1 - Cdh2 |  | 0.87 *** | -1.29 * | -1.25 . | -0.53 ** | -0.52 *** | 1.04 *** |
| Hypothalamus | Oligodendrocytes Mog Plp1 - Mal -> Oligodendrocytes Mag Plp1 - Opalin | Plp1 | 1.06 * | -0.47 *** | 1.29 *** | -1.79 *** | -0.1 . | 0.05 . |
|  | Oligodendrocytes Mog Plp1 - Plp1 -> OPC Pdgrfra Plp1 - Lhfp13 |  | -1.49 ** | 2.42 *** | 4.16 *** | -0.18 . | -0.01 . | 0.13 . |
|  | OPC Pdgrfra Gpr17 - S100a1 -> Oligodendrocytes Mag Gpr17 - Frmd4a | Gpr17 | 2.89 *** | -4.67 *** | -1.68 * | 0 . | -0.62 . | -0.24 . |
|  | Oligodendrocytes Olig2 Gpr17 - Tnr -> Oligodendrocytes Mag Gpr17 - Frmd4a |  | 2.12 *** | -2.84 *** | -1.53 . | 0.14 . | -0.48 . | -0.19 . |
|  | Oligodendrocytes Mag Gpr17 - Frmd4a -> OPC Pdgrfra Gpr17 - S100a1 |  | -2.89 *** | 4.67 *** | 1.68 * | 0 . | 0.62 . | 0.24 . |
|  | Oligodendrocytes Mag Gpr17 - Frmd4a -> Oligodendrocytes Olig2 Gpr17 - Tnr |  | -2.12 *** | 2.84 *** | 1.53 . | -0.14 . | 0.48 . | 0.19 . |
| Orbitofrontal cortex | Astrocytes Slc1a3 Car2 - Glul -> Glutamatergic Cck Car2 - Aplp1 | Car2 | 2.77 *** | -0.86 *** | -0.22 . | -0.08 . | -0.19 . | -0.07 . |
|  | Astrocytes Slc1a3 Car2 - Glul -> Oligodendrocytes Mog Car2 - Plp1 |  | 2.43 *** | -1.49 *** | -0.4 . | -0.35 . | -0.11 . | -0.1 . |
|  | Astrocytes Slc1a3 Car2 - Cd81 -> Oligodendrocytes Mog Car2 - Plp1 |  | 2.36 *** | -1.16 *** | -0.72 *** | -0.16 . | 0.12 . | 0.01 . |
| Organoids | NSCs SOX2 CDC20 - CCNB2 -><br>NSCs SOX2 CDC20 - CA2 | CDC20 | -3.94 *** | -2.59 ** | -2.04 *** | -0.42 . | -0.27 * | 0 . |
|  | NSCs SOX2 CDC20 - CCNB1 -><br>NSCs SOX2 CDC20 - CA2 |  | -3.72 *** | -2.29 * | -2.12 *** | -0.27 . | -0.24 * | 0 . |
|  | NSCs SOX2 CDC20 - MKI67 -><br>NSCs SOX2 CDC20 - CA2 |  | -3.71 *** | -2.75 *** | -2.07 *** | -0.3 . | -0.2 . | 0 . |
|  | NSCs SOX2 CDC20 - CA2 -><br>NSCs SOX2 CDC20 - NUF2 |  | 3.67 *** | 2.33 . | 2.57 *** | 0.59 . | 0.27 . | 0 . |
|  | NSCs SOX2 CDC20 - CA2 -><br>NSCs SOX2 CDC20 - CDC20 |  | 3.96 *** | 3.01 *** | 1.78 ** | 0.24 . | 0.27 * | 0 . |
| Striatum | GABAergic Sst Npy - Npy -><br>GABAergic Sst Npy - Sst | Npy | 0.36 *** | 7.47 *** | 1.57 * | 0.89 *** | 1.29 *** | 0 . |
|  | Cholinergic Tacr1 Npy - Tacr1 -><br>GABAergic Sst Npy - Sst |  | -0.06 . | 7.4 *** | 1.7 . | 0.69 ** | 3.16 *** | 0 . |

## Discussion

Data generated in massive life science experiments are analytically challenging since they contain many variables resulting in high volumes of data. A particular example of large variability is scRNAseq data containing a high number of individual transcripts within a high number of single-cell transcriptomes. Such data complexity demands a multistage analysis, mainly including dimensionality reduction, clustering, and selection of significantly deregulated features between clusters, and requires defining multiple parameters, such as PC, neighbors, distance, etc. Countless parameter settings result in the generation of a large number of potential false positive and false negative results leading to the overinterpretation of experiments.

There are a variety of scRNAseq analytical tools containing straightforward analytical methods, for instance, dimensional reduction by principal components followed by the secondary dimensional reduction based on UMAP or t-SNE used for cluster visualization. The correct principal components are selected based on the visual presentation of the percent of explained variance and JackStraw analysis of the PC significance. Adding even more principal components to explain more data variance can result in a more positive influence on results; however, such an approach is also associated with an increase in dimensionality, which can negatively impact the results. Therefore, manipulating the number of PCs can not explain the whole heterogeneity of cell populations but may only improve it. However, the experimenter is still unsure if added or subtracted even statistically relevant PCs reveal authentic or artificial biological final results.

Such caveats about the danger of omitting some specific components with relevant information are often debated within the single-cell-specialized research community (https://satijalab.org/seurat/articles/pbmc3k_tutorial.html or https://jkubis96.github.io/Supplement_JSEQ_scRNAseq/supplementary_data/Enrichment/seurat_community.bmp). Therefore, the decision to select relevant PCs is crucial; unfortunately, too much depends on user decision which may lead to misinterpretation. Such initial reduction of dimensionality is essential and influences further steps of cell clustering. In addition, subsequent steps mentioned above, namely KNN and SNN, require even more adjustable and obligatory parameters. Using such methods, we can quickly obtain the general cell diversity of the tissues and visualize dominant cell populations. The heterogeneity offered by such an approach is suitable for many applications however does not allow for efficient and reliable detection of intermediate and instant cell stages, trajectories, and cell fates which is critical for dynamic processes such as development and disease progression.

Addressing all the aspects needs a new approach for the bioinformatic processing of single-cell data and for establishing high-resolution tissue profiles such as the brain. However, several conditions need to be fulfilled for a successful approach. First, the data flow between applications that starts with reads mapping, repairing, and determination of basic clusters, etc., needs to be always uniform. Second, the applications need interoperability between each other to pass the data according to the hand-to-hand principle with equal parameters, which means automatic and decisive data handling between all applications. The third requirement is the selection and preservation of high-quality data, and if the data are of non-sufficient quality, they should first undergo the improvement of quality by our repairing algorithms. The aim is to retrieve as much information from the dataset as possible and represent uniform pre-processed quality for feeding the CSSG algorithm to later explain cell population heterogeneity with high resolution.

Such data processing can decrease the possibility that the discovered rare populations are artificial and exist without the support of their lineage and should provide the experimenter with needed confidence. The approach eliminates manual parameter change and runs repeated calculations in search of convincing dimensional reduction for a particular rare population. Moreover, the cell number of a particular subtype is additionally subjected to quality control by binominal test; therefore, the probability of inclusion of artificial subtypes is minimized.

The most powerful feature of the CSSG algorithm is its focus on the real biological values of the differentiated occurrence of genes selected as marker genes for a given cluster. In subsequent iterations of operations on the matrices, the algorithm selects the best combination of marker genes, the presence of which is best described by the heterogeneity of the cluster, filling the gaps between maker genes. Thanks to this, we are able to identify the best combination of genes that explain the internal diversity of the cluster. The obtained combinations of genes constitute a minimal element explaining the real biological heterogeneity resulting from the constantly changing functions of cells derived from the same lineage during their life, which often results from the variability in the expression of individual single genes. Importantly, every dataset has its individual profile of heterogeneity which is determined by sequencing depth, library preparation, cell lineage in a sample, and sample processing. Therefore it is critical to analyze the datasets from other sources independently in order to obtain only biologically relevant differences and avoid experiment-related bias.

Thanks to this approach, we established 620 cell subtypes derived from 239 base cell subclasses across the different parts of the mouse brain and in the human brain organoids. Moreover, based on the CSSG-selected markers presented in obtained subtypes, we could notice that they differed in the development stages, maturation status, and signaling. Therefore the results were used to demonstrate the pipeline advantage to establish exact trajectories of cell population inside and between cell clusters.

For instance, the oligodendrocytes are a powerful model of brain cell differentiation, presenting many intermediate cell stages. We defined the model of oligodendrocyte lineage by the collective screening of many works describing precursors and mature cells. The works show oligodendrocyte lineages of OPC, COP, preMO, NFO, MFO, and MOL (37–41). In brief, the markers for OPCs are Vcan (37), C1ql1 (38), and Cspg5, and S100a1 (37, 39) and Xylt1, important in the pathway (42) leading to the synthesis of chondroitin sulfate proteoglycans (CSPG) involved in OPC differentiation towards myelin-producing cells (40). Moreover, in the literature, the COP vs. OPC cells show higher expression of Tns3 and Vcan, which are marker parts of model (37). The preMO as intermediate cells was also proposed as the cell stage before NFO oligodendrocytes, and in addition, the preMO stage starts to express Mag (41). Our literature model points on Gpr17 gene as related to preMO (41) and to the early development of oligodendrocytes (23, 37, 41). In addition, we have selected Tnr gene as another preMO marker candidate. The Tnr is involved in oligodendrocytes differentiation but inhibits myelin membrane formation (43). The next form is the NFO presenting Frmd4a and Fyn genes (37), the St18, which is needed for oligodendrocytes formation (44). The MOL phase express Anl (37), Mbp (45), Prr5l (45), and co-expression of Car2 with Plp1 is related to this cell stage (46). Finally, the expression of Mog (23), Opalin, Mal, and Plp1 (37) genes responsible for myelin formation are widely expressed in MFOL and MOL.

Trajectories of cell differentiation can be determined based on analyzing the subtypes expressing the dominant genes selected by CSSG, which are the molecular switches of trajectories. The relations between the subtypes can now be determined by hierarchical clustering, which builds the cell population trajectories. The above paradigm of CSSG and hierarchical clustering was used for the oligodendrocyte molecular switches and subpopulation relations. We noticed that the paradigm results showed the oligodendrocyte’s decreasing maturity along the hierarchy in the lineage tree. To assign the names of the lineage stages, we used the literature-based model of oligodendrocyte differentiation described above and confronted with markers forming names of our cell population of subclass and subtypes (see the cell population naming rules). Such a complex model of trajectories was never shown previously with such resolution in one experiment using complex brain tissue.

In a similar approach as trajectories and as another pipeline validation step, we traced the relations between cell populations (types, subclasses, and subtypes) specifically expressing Alzheimer’s disease genes such as Apoe, App, Mapt, Psen1, Psen2, and Trem2 in mouse brain and human brain organoids (Fig. 5, 6, 7 and supplement 1.3.1).

Increased expression of the Apoe gene is characteristic for astrocyte, microglial endothelial, and pericyte cells (47–50). The expression of Apoe is also present in OPCs and oligodendrocytes, where it is involved in the migration and maturation of OPCs and oligodendrocytes and general brain remyelination (51). In neurons, the Apoe silencing results in the downregulation of proliferation in favor of differentiation, followed by increased excitatory current and release of neurotransmitters (52). Using the CSSG/JSEQ, we demonstrated high expression of Apoe in subtypes of non-neuronal cells and in several glutamatergic subtypes.

App-Apoe protein product interaction regulates Aβ aggregation and clearance and may influence the development of plaques and tau pathology (53). In our results, the co-expression of the genes is characteristic for some vascular (endothelial, pericytes, fibroblast), immune, astrocyte, glutamatergic, and NSC subtypes. Our analysis can pinpoint the exact subtypes where such an interaction may potentially occur and become an accurate AD therapy target.

In general, the expression pattern of AD-involved genes is already complicated in cell types; therefore, it is impossible to explain cellular disease complexity without splitting the proposed cell populations into subtypes. For instance, Apoe, Trem2, and others are traditionally regarded as glial markers, but both markers’ expression is also found in several subtypes of neuronal cells. For the orbitofrontal subclass (initial cluster), we obtained four subtypes (Fig. 8) that belong to up to three different cell types, including astrocytes, oligodendrocytes, and glutamatergic cells. At the highest level, two hierarchical clusters in our graph contain two astrocytic populations with different markers. We observed astrocytes expressing Cd81, a critical regulator of neuronal-induced astrocytic differentiation (54), next to the classical astrocyte subtype expressing Glul (55). Next, our hierarchy revealed the glutamatergic cell subtype, followed by the oligodendrocyte subtype. Such hierarchical assignment of subtypes generated by the CSSG/JSEQ approach is very interesting since some populations of astrocytes may acquire new functions. Some astrocytes, namely their intermediate populations, may release glutamate (56) or may convert to oligodendrocytes (57, 58) in response to injury or disease stimuli. Another candidate for intermediate cellular subtypes can be identified in the subclass of macrophages in the entorhinal cortex, where the particular subtype of macrophages expresses Cdh2 gene. We hypothesized that the population might be in the intermediate subtype stage characteristic of glutamatergic cells. Also, in response to inflammation, the microglial can start the production of glutamate (59). The CSSG/JSEQ isolated cellular intermediate stages show marked patterns of genes expression causative to AD. The analysis can point at new cellular targets and the relation of AD gene expression to search for new therapies.

## Conclusion

Single-cell RNA sequencing generates high volumes of data demanding complicated multi-step bioinformatics analysis and specialized applications. Existing solutions are not reliably applicable to discover the high level of tissue heterogeneity, cellular lineages during development, and differentiation with many intermediate rare populations. Therefore we have generated a new algorithmic strategy and pipeline for rare cell population discovery which we called CSSG/JSEQ. The CSSG algorithm processes scRNAseq data expressed as binary matrix data about genes occurrence in each cell inside the cluster in an iterative and parallel way. The action of the CSSG is complemented with a new naming algorithm that uses a cascade of CSSG results and the metadata of marker genes to generate names of cell classes, subclasses, and subtypes. We conclude that the CSSG and naming approach allows for tracing cell populations with high resolution, finding relations between them to data mine the hierarchy between cell populations, and establishing developmental, differentiation, and pathogenic trajectory. The important advantage of our pipeline is that it is able to take a “snapshot” of a current physiological or pathological stage of the investigated organ or a cell population which is particularly important for complicated tissues such as the brain.

## Supplementary Data

Supplementary data (supplement 1.1, supplement 1.2, and supplement 1.3 [1.3.1 & 1.3.2]) have been combined into one HTML file containing extended results for all brain regions and human brain organoids. The supplement is available at the following link for convenience:

[https://jkubis96.github.io/Supplement_JSEQ_scRNAseq/].

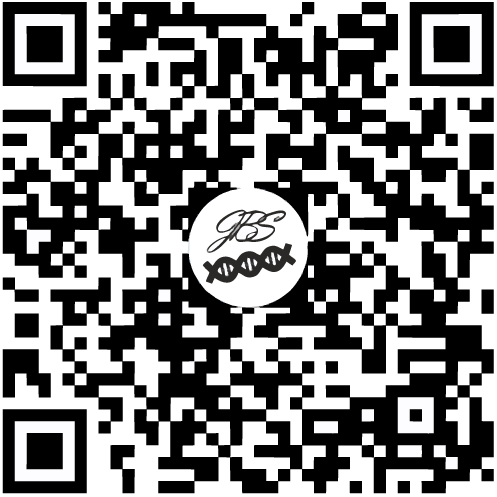

## Acknowledgment

We sincerely thank the Broad Institute for generating their open-access single-cell data portal (https://singlecell.broadinstitute.org/single_cell). Sustaining and updating the comprehensive single-cell dataset freely available to researchers from the community greatly accelerates the development of single-cell research and dedicated bioinformatics solutions like creating and testing our CSSG/JSEQ algorithms and pipeline.

